# Flat Electrode Contacts for Peripheral Nerve Stimulation

**DOI:** 10.1101/593467

**Authors:** Jesse E Bucksot, Andrew J Wells, Kimiya C Rahebi, Vishnoukumaar Sivaji, Mario Romero-Ortega, Michael P Kilgard, Robert L Rennaker, Seth A Hays

**Affiliations:** The University of Texas at Dallas, Erik Jonsson School of Engineering and Computer Science, Richardson, Texas, U.S.; Texas Biomedical Device Center, Richardson, Texas, U.S.; The University of Texas at Dallas, School of Behavioral Brain Sciences, Richardson, Texas, U.S.

## Abstract

The majority of available systems for nerve stimulation use circumferential stimulation electrodes inside an insulating cuff, which produce largely uniform current density within the nerve. Flat stimulation electrodes that contact only one side of the nerve may provide advantages including simpler implantation, ease of production, and more resistance to mechanical failure. However, it is possible that the flat configuration will yield inefficient fiber recruitment due to a less uniform current distribution within the nerve. Here we tested the hypothesis that flat electrodes will require higher current amplitude to achieve effective stimulation than circumferential designs. Computational modeling and in vivo experiments were performed to evaluate fiber recruitment in different nerves and different species using a variety of electrode designs. Initial results demonstrated similar fiber recruitment in the rat vagus and sciatic nerves with a standard circumferential cuff electrode and a cuff electrode modified to approximate a flat configuration. Follow up experiments comparing true flat electrodes to circumferential electrodes on the rabbit sciatic nerve confirmed that fiber recruitment was equivalent between the two designs. These findings demonstrate that flat electrodes represent a viable design for nerve stimulation that may provide advantages over the current circumferential designs for applications in which the goal is uniform activation of the nerve.

## Introduction

Peripheral nerve stimulation has emerged as an effective treatment for a variety of disorders. Vagus nerve stimulation (VNS) is one of the most widely used peripheral nerve stimulation strategies and has been employed in over 70,000 patients for control of epilepsy (1). Recent clinical studies demonstrate the potential of VNS for treatment of other neurological disorders, including stroke (2), tinnitus (3), and headache (4). Other emerging nerve stimulation therapies include tibial nerve stimulation for bladder control (5), sacral nerve stimulation for constipation (6), occipital nerve stimulation for migraines (7), and hypoglossal nerve stimulation for sleep apnea (8). Given the broad potential applications for neurological disorders, there is a great deal of interest in identifying optimal stimulation strategies to maximize benefits in patients (9).

Implanted cuff electrodes are the gold-standard method for nerve stimulation. The majority of recent developments have been focused on selective stimulation of certain regions or fascicles within the nerve (10–12). Such designs could potentially eliminate side-effects that arise due to stimulation of off-target fibers and could allow for more precise stimulation of target fibers. However, in other applications including VNS, the primary goal based on existing evidence is to generate uniform activation of each region of the nerve. This is typically done with circumferential or helical electrodes that cover the majority of the circumference of the nerve. These designs provide largely uniform stimulation within the nerve yielding a steep recruitment curve and ensuring that any desired fibers are activated with minimal current.

Alternative designs may provide better avenues for nerve stimulation and obviate common issues such as a high rate of lead breakage (13). A flat configuration with electrode contacts on one side of the nerve could be built with a simpler and more compact design that would facilitate implantation, provide greater resistance to mechanical failure, and reduce cost of production. However, this electrode geometry provides contact with only a portion of the circumference of the nerve, which is likely to produce non-uniform stimulation. This would yield increased activation of axons near the contacts and reduced activation of distant axons. The resulting polarity would lead to a lower threshold and higher point of saturation and thus a less steep recruitment curve. Whether the magnitude of this effect would substantially influence efficacy is not known. A direct comparison of flat and circumferential cuff electrodes is needed to determine if flat contacts represent a practical alternative for nerve stimulation.

## Materials and Methods

### Computational Model

A 3D model was created in Comsol (COMSOL Multiphysics® Version 5.3) consisting of a nerve with a single fascicle, perineurium, epineurium, two platinum contacts, an insulating cuff, and ambient medium, similar to previous studies (14,15). In a subset of models, a multi-fascicle nerve containing five fascicles was used (Fig. 9). The nerve had a diameter of either 0.9 mm for the rat sciatic, 0.4 mm for the rat vagus, or 3 mm for the rabbit sciatic (16–18). Perineurium thickness was set to 3% of the fascicle diameter (19). Epineurium thickness was set to 0.13 mm for the rat sciatic, 0.1 mm for the rat vagus, and 0.43 mm for the rabbit sciatic (20–22). To investigate the effect of nerve size, the rabbit sciatic model was scaled from 4 times smaller to 1.5 times larger. For both rat nerves, the insulating cuffs had an inner diameter of 1 mm and outer diameter of 2 mm. For the rabbit sciatic, insulating cuffs had an inner diameter of 3.02 mm and outer diameter of 5.2 mm. Platinum contacts had a thickness of 0.01 mm. Flat contacts used in the rabbit model had a width of 2 mm and length of 1.5 mm in the axial direction. The cross-sectional area of the nerve and of the cuff lumen was matched between the circumferential and flat electrode models by increasing the inner diameter of the flat cuff by 8.67%. The nerve was modified to take on the shape of the flat cuff (Fig. 6) (23). Helical electrodes with a width of 0.7mm and thickness of 0.01mm had a pitch of 2 mm and completed a 270° arc. The insulation had the same pitch and completed 2.5 turns. The width of the insulation was 1.4 mm and the thickness was 0.9 mm. The empty space in all models was filled with an ambient medium with conductance varying from saline (2 S/m) to fat (0.04 S/m). For the rat models, ambient mediums were 20 mm in length and 4 mm in diameter. For the rabbit models, they were 120 mm in length and 40 mm in diameter. The outer boundaries of all models were grounded. A 1 mA positive current was applied on one contact, and a 1 mA negative current on the other. Due to the model being purely resistive, the voltage field only needed to be solved for a single current amplitude. Electrical properties for each material were based on field standards and can be found in table S1 (24–27).

Once the Comsol model was solved, the voltage distribution inside the fascicle was exported and read into a NEURON model consisting of 500 parallel axons uniformly distributed throughout the fascicle. The multi-fascicle nerve had 100 axons in each of the five fascicles. Axons were designed using the model created by McIntyre, Richardson, and Grill (28). All electrical parameters were identical in this study, but the geometric parameters were interpolated using either a 1^st^ or 2^nd^ order polynomial. All fitted functions can be found in table S2. Each fiber was set to the length of the corresponding Comsol model, either 20 mm or 120 mm. Diameters were taken from a normal distribution meant to represent A-fibers (rat sciatic: 6.87 ± 3.02 µm, rat vagus: 2.5 ± 0.75 µm, rabbit sciatic: 8.85 ± 3.1 µm) (29,30). Rat vagus fiber diameters were estimated based on the conduction velocity of the fibers mediating the Hering-Breuer reflex (31–34). In both sciatic models, fibers with a diameter less than 2 µm were recreated until their diameter was greater than 2 µm. In the vagus model, the same technique was applied but with a cutoff of 1 µm.

After a 0.5 ms delay to ensure all axons had reached a steady baseline, a biphasic pulse of varying current amplitude was applied to the NEURON model. The voltage field calculated in Comsol was linearly scaled to the specified current and applied for 0.1 ms, and then the inverse was applied immediately after for another 0.1 ms. Voltage traces from nodes at the proximal end of the axon were recorded and used to determine whether that axon was activated at the given current amplitude. The activation data was used to create dose-response curves showing the percentage of axons activated as a function of current amplitude.

### Animal Experiments

All handling, housing, stimulation, and surgical procedures were approved by The University of Texas at Dallas Institutional Animal Care and Use Committee. Twelve Sprague Dawley female rats (Charles River, 3 to 6 months old, 250 to 500 g) were housed in a 12:12 h reverse light-dark cycle. Six rats were used for sciatic experiments, and six rats were used for vagus experiments. Four New Zealand white male rabbits (Charles River, 3 to 6 months old, 2 to 4 kg) were housed in a 12:12 h light-dark cycle. All four rabbits were used for sciatic experiments.

### Electrodes

Rat experiments were performed using custom-made cuff electrodes. All cuff electrodes were hand-made according to standard procedures (35). The cuffs were insulated with 3 to 6 mm sections of polyurethane tubing with an inner diameter of 1 mm and outer diameter of 2 mm. Electrodes were multi-stranded platinum-iridium wire with a diameter of 0.01 mm. For the circumferential cuff electrode, platinum-iridium wires covered a 270° arc inside the cuff. To approximate a flat electrode, partial contacts were used which only covered a 60° arc. Additionally, an intermediate electrode was tested with a 120° arc. All electrode impedances were measured in saline before testing to ensure proper construction.

Rabbit experiments were performed using both custom-made circumferential electrodes and manufactured flat electrodes. The circumferential electrodes were made using the same materials and protocol as the rat cuff electrodes but sized to accommodate the larger rabbit sciatic nerve (3 mm inner diameter, 4.5 mm outer diameter, 270° arc). Flat electrodes consisted of PCBs connected to two rectangular platinum contacts (36). All on-board components were encapsulated and hermetically sealed in glass. Current controlled stimulation was delivered with this device using an on-board microcontroller with a digital to analog converter (DAC). Analog output from the DAC was amplified by an op-amp with a maximum current of 1.2 mA. The rectangular contacts were attached to the surface of the glass and connected to the PCB through hermetic through glass vias. A 9-turn 3-layer coil was used as an antenna for power reception and communication. A silicone sleeve was fitted around the device to serve as an insulating cuff.

### Rat Sciatic Nerve Stimulation

Rats were anesthetized using ketamine hydrochloride (80 mg/kg, intraperitoneal (IP) injection) and xylazine (10 mg/kg, IP) and given supplemental doses as needed. Once the surgical site was shaved, an incision was made on the skin directly above the biceps femoris (16,37). The sciatic nerve was exposed by dissecting under the biceps femoris. The gastrocnemius muscle was separated from skin and surrounding tissue. Cuff electrodes were then placed on the sciatic nerve with leads connected to an isolated programmable stimulator (Model 4100; A-M Systems™; Sequim, WA). The nerve was left in place underneath the biceps femoris and the cavity was kept full of saline at all times to ensure that the cuff would be operating in a uniform medium with conductance similar to tissue. The Achilles tendon was severed at the ankle and affixed to a force transducer using nylon sutures. The foot was clamped and secured to a stereotaxic frame to prevent movement of the leg during stimulation and to isolate recordings from the gastrocnemius muscle.

Stimulation was delivered through the A-M Systems™ Model 4100. Voltage traces were recorded using a digital oscilloscope (PicoScope® 2204A; Pico Technology; Tyler, TX). The force of muscle contraction was recorded through a force transducer (2kg EBB Load Cell; Transducer Techniques; Temecula, CA) which was connected to an analog channel on an Arduino® Mega 2560. All components were integrated using MATLAB®. Data was sampled at 10 Hz.

Stimulation consisted of 0.5 second trains of biphasic pulses (100 µs pulse width) at 30 Hz with varying current amplitudes ranging from 20-800 µA. Stimulation intensities were randomly interleaved. Values for current were manually set in each experiment to ensure that the range of values included the entire dose-response curve. Stimulation was delivered every 15 seconds and each parameter was repeated in triplicate.

### Rat Vagus Nerve Stimulation

Rats were anesthetized using ketamine hydrochloride (80 mg/kg, IP) and xylazine (10 mg/kg, IP) and given supplemental doses as needed. An incision and blunt dissection of the muscles in the neck exposed the left cervical vagus nerve, according to standard procedures (38–40). The nerve was placed into the cuff electrode, and leads from the electrode were connected to the programmable stimulator. The cavity was kept full of saline at all times. To assess activation of the vagus nerve, blood oxygen saturation (SpO2) was recorded using a pulse-oximeter (Starr Life Sciences™, MouseOx Plus®) as previously described (32). Data was read into MATLAB® using a Starr Link Plus™ with the outputs connected to analog channels on the Arduino®. Data was sampled at 10 Hz and filtered with a 10 sample moving average filter.

Stimulation consisted of 5 second trains of biphasic pulses (100 µs pulse width) at 30 Hz with varying current amplitudes ranging from 50-2500 µA. Values for current were randomly interleaved.

Stimulation was delivered every 60 seconds, but was delayed if needed to allow the oxygen saturation to return to baseline. Each parameter was repeated twice.

#### Rabbit Sciatic Nerve Stimulation

Both hind legs of the rabbit were shaved over the incision site the day before surgery. Anesthesia was induced with 3% inhaled isoflurane at 3 L/min. A single intraperitoneal injection of ketamine hydrochloride (35 mg/kg) and xylazine (5 mg/kg) was given after induction. Isoflurane was maintained throughout the experiment. Eye ointment was applied to both eyes to prevent drying. Rectal temperature and breathing were monitored throughout the procedure. The incision sites were cleaned with 70% ethanol, followed by povidone-iodine, followed again by 70% ethanol. An incision site was made along the axis of the femur. The sciatic nerve was exposed with blunt dissection to separate the biceps femoris and quadriceps femoris muscles. Alm retractors were placed to allow cuff implantation. After placing the cuff around the nerve, the retractors were withdrawn.

Stimulation consisted of 0.5 second trains of biphasic pulses (100 µs pulse width) at 10 Hz with varying current amplitudes ranging from 20-1600 µA. Stimulation using the circumferential cuff electrode was delivered using the same system described above for the rat sciatic. Stimulation with the glass-encapsulated electrode was delivered directly from the PCB. The on board stimulation circuit had a resolution of 33µA, which was too large to accurately fit sigmoid functions to the fiber recruitment curve in most cases. Values for current were randomly interleaved. Stimulation was delivered every 5 seconds and each parameter was repeated in triplicate. Data was sampled at 500 Hz using the same load cell collection system described above.

### Analysis and Statistical Comparisons

All responses were normalized to the maximum response recorded in each subject. Raw, non-normalized responses are included in the supplementary information (Fig. S4-S6). Dose-response curves were fitted with a sigmoid function (Fig. 1c). Restrictions were placed on the fitted curve such that the point at 1% of Y_max_ could not be at a negative current intensity. For each curve, the slope was calculated at the midpoint of the fitted function. The threshold was determined by finding the lowest current amplitude that always resulted in a change in the signal of force or SpO2 greater than 3x the standard deviation of the preceding 1 second of signal for muscle activation or 10 seconds of signal for SpO2. The saturation point was determined by finding the lowest current value that produced a change in the signal greater than 90% of the mean of the top 50% of the curve. The dynamic range was calculated as the saturation point minus the current value one step below the threshold. All analyses were verified by a blinded experimenter.

**Fig 1.**
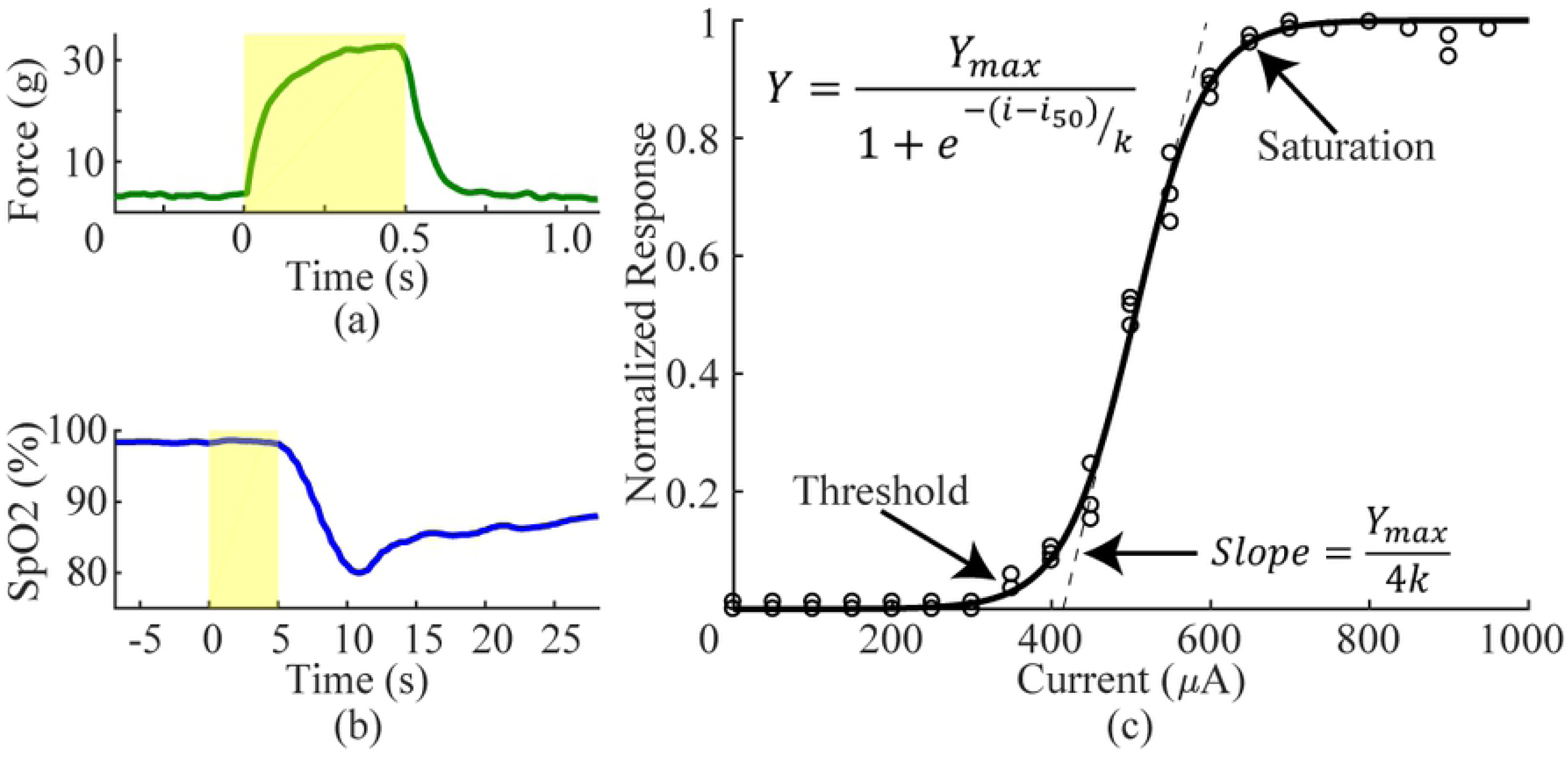
Analysis of fiber recruitment. a) To assess sciatic nerve recruitment, we measured force of hindlimb muscle contraction in response to a range of stimulation intensities. Shaded region represents stimulation at 30Hz for 0.5 seconds. b) To measure vagus nerve recruitment, we measured reductions in SpO2. Shaded region represents stimulation at 30Hz for 5 seconds. c) An example recruitment curve with a fitted sigmoid function and all outcome measures identified.

Data reported in the text and figures represent mean ± standard error of the mean (SEM). The sample size shown in each figure is equal to the total number of experiments performed with each electrode design, not the number of animals. Thresholds, saturation points, dynamic ranges, and slope of each electrode design were compared on the rat sciatic using a one-way ANOVA after confirming equal variance with a Bartlett test. Individual comparisons were made using post-hoc Tukey-Kramer tests. For the rat vagus and rabbit sciatic, the variance of each metric was compared with a two-sample F-test and then the data were compared using a two tailed two-sample t-test with either equal or unequal variance depending on the F-test. The statistical test used for each comparison is noted in the text. All calculations were performed in MATLAB.

## Results

### One-sided and circumferential electrodes provide equivalent recruitment of rat sciatic nerve

Flat electrode contacts that do not surround the entire nerve may yield less efficient fiber recruitment than circumferential electrode contacts. We tested recruitment efficacy using computational modeling and *in vivo* experiments on the rat sciatic nerve. To represent flat electrodes, we used a modified circumferential electrode that only provided 60° of coverage compared to the standard 270° (Fig. 3b). Fiber recruitment functions were created using the 60° and 270° designs as well as an intermediate 120° design.

#### Model

We used computational modeling to evaluate fiber recruitment using multiple electrode designs with different values for angle of coverage, contact spacing, cuff overhang, and cuff inner diameter. Reducing the angle of coverage had a small effect on recruitment (Fig. 2d). The smallest angle (30°) required 105.2 µA to recruit 5% of fibers (*i*_5%_) whereas the standard angle (270°) required 143.4 µA. To recruit 95% of fibers (*i*_95%_), the smallest angle required 311.7 µA and the standard angle required 296.0 µA.

**Fig 2.**
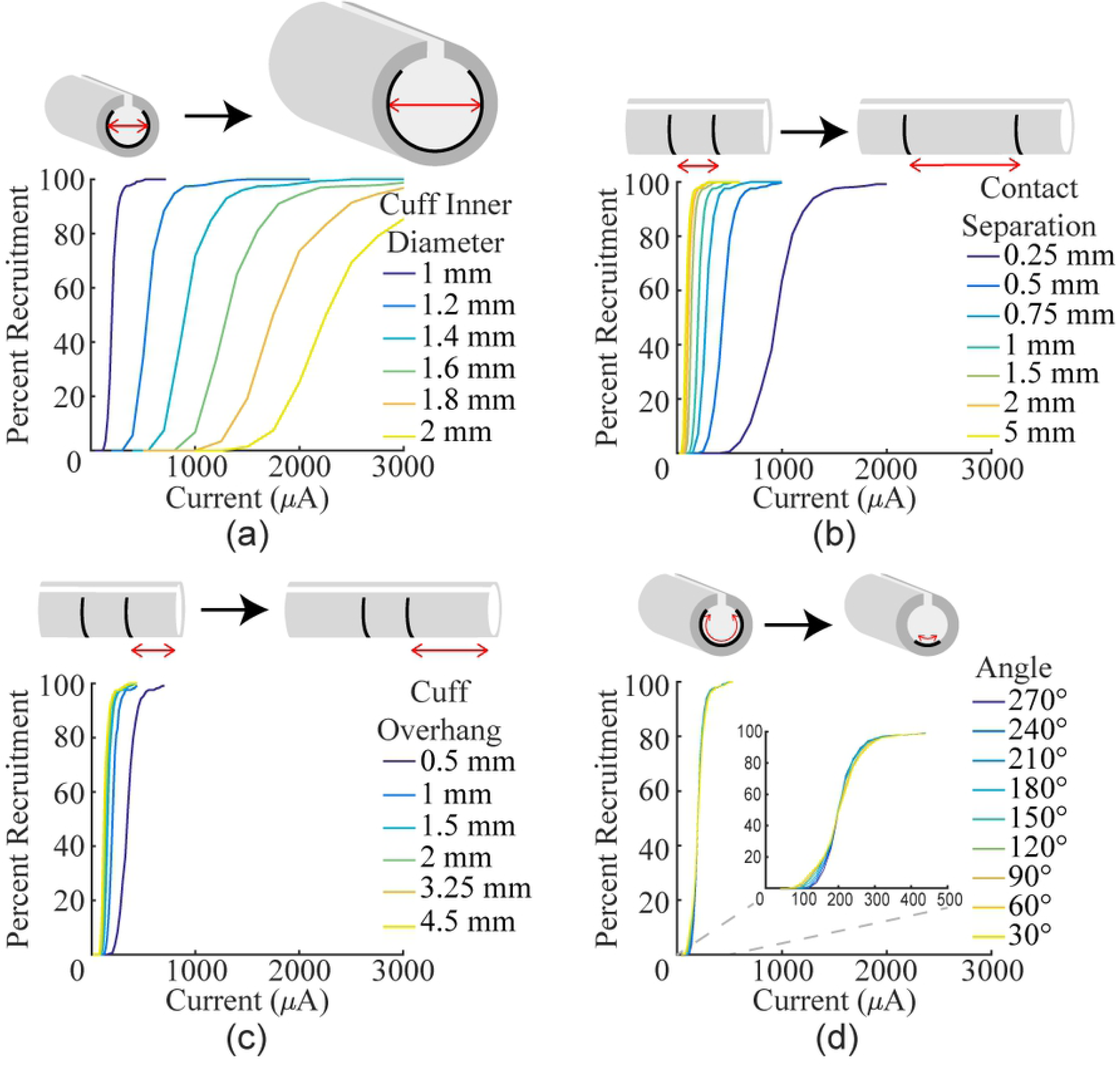
Modeling the effect of various cuff electrode parameters on recruitment of rat sciatic nerve. Recruitment curves were generated with different values for several design parameters. a) Increasing the inner diameter of the cuff (1 mm contact separation, 1 mm cuff overhang, 270°) drastically reduces recruitment. b) Increasing the distance between the two stimulating contacts (1 mm cuff inner diameter, 1 mm cuff overhang, 270°) increases recruitment. c) Increasing the amount of cuff overhang (1 mm cuff inner diameter, 1 mm contact separation, 270°) increases recruitment. d) Reducing the angle of coverage (1 mm cuff inner diameter, 1 mm contact separation, 1 mm cuff overhang) has minimal effect on recruitment.

Unlike angle of coverage, the other three variables strongly influenced recruitment. With a standard 270° arc, increasing the inner diameter of the cuff had the strongest effect on recruitment, greatly increasing both the threshold and saturation current (Fig. 2a; 1 mm: *i*_5%_=143.4 µA, *i*_95%_=296.0 µA; 1.2 mm: *i*_5%_=378.6 µA, *i*_95%_=807.7 µA). Increasing the distance between the contacts lowered both the threshold and saturation current (Fig. 2b; 0.25 mm: *i*_5%_=332.3 µA, *i*_95%_=697.3 µA; 5 mm: *i*_5%_=41.8 µA, *i*_95%_=119.2 µA). Increasing the amount of cuff overhang lowered both the threshold and saturation current (Fig. 2c; 0.5 mm: *i*_5%_=232.3 µA, *i*_95%_=490.0 µA; 4.5 mm: *i*_5%_=88.4 µA, *i*_95%_=202.1 µA). Compared to the impact of the three other variables, angle of coverage was the least important factor, suggesting that it is not a critical factor in electrode design and flat electrodes would achieve saturation at similar current amplitudes to circumferential electrodes. Varying the cuff inner diameter, contact separation, and cuff overhang on a 60° electrode demonstrated that each variable affects an electrode with a shorter arc just as it does a standard electrode (Fig. S1).

#### Empirical

To confirm modeling predictions, we evaluated nerve recruitment in the rat sciatic nerve using the 60°, 120°, and 270° electrodes. In vivo data closely resembled data derived from the model, with flat and circumferential contacts demonstrating comparable fiber recruitment. No significant differences were found between recruitment thresholds for any of the electrode configurations (Fig. 3d; 60°: 131.7 ± 14.2 µA, 120°: 134.4 ± 18.3 µA, 270°: 135.0 ± 15.4 µA; Bartlett’s test, χ2(0.05, 2)=0.110, p=0.94639, one-way ANOVA, F(2, 29)=0.01, p=0.986). Additionally, ANOVA did not reveal differences in saturation current, dynamic range, or slope between the electrode designs (Saturation: Fig. 3e, 60°: 205.0 ± 23.6 µA, 120°: 195.6 ± 27.2 µA, 270°: 176.4 ± 20.0 µA, Bartlett’s test, χ2(0.05, 2)=0.448, p=0.799, one-way ANOVA, F(2, 29)=0.41, p=0.569; Dynamic range: Fig. 3f, 60°: 93.3 ± 13.6 µA, 120°: 80.0 ± 10.5 µA, 270°: 60.0 ± 7.6 µA, Bartlett’s test, χ2(0.05, 2)=4.310, p=0.116, one-way ANOVA, F(2, 29)=2.41, p=0.0647; Slope: Fig. 3g, 60°: 2.21 ± 0.31 %Force/µA, 120°: 2.63 ± 0.41 %Force/µA, 270°: 3.71 ± 0.63 %Force/µA, Bartlett’s test, χ2(0.05, 2)=3.195, p=0.202, one-way ANOVA, F(2, 29)=3.03, p=0.065). These results confirm that the one-sided electrodes and circumferential electrodes yield equivalent nerve recruitment across a range of stimulation intensities.

**Fig 3.**
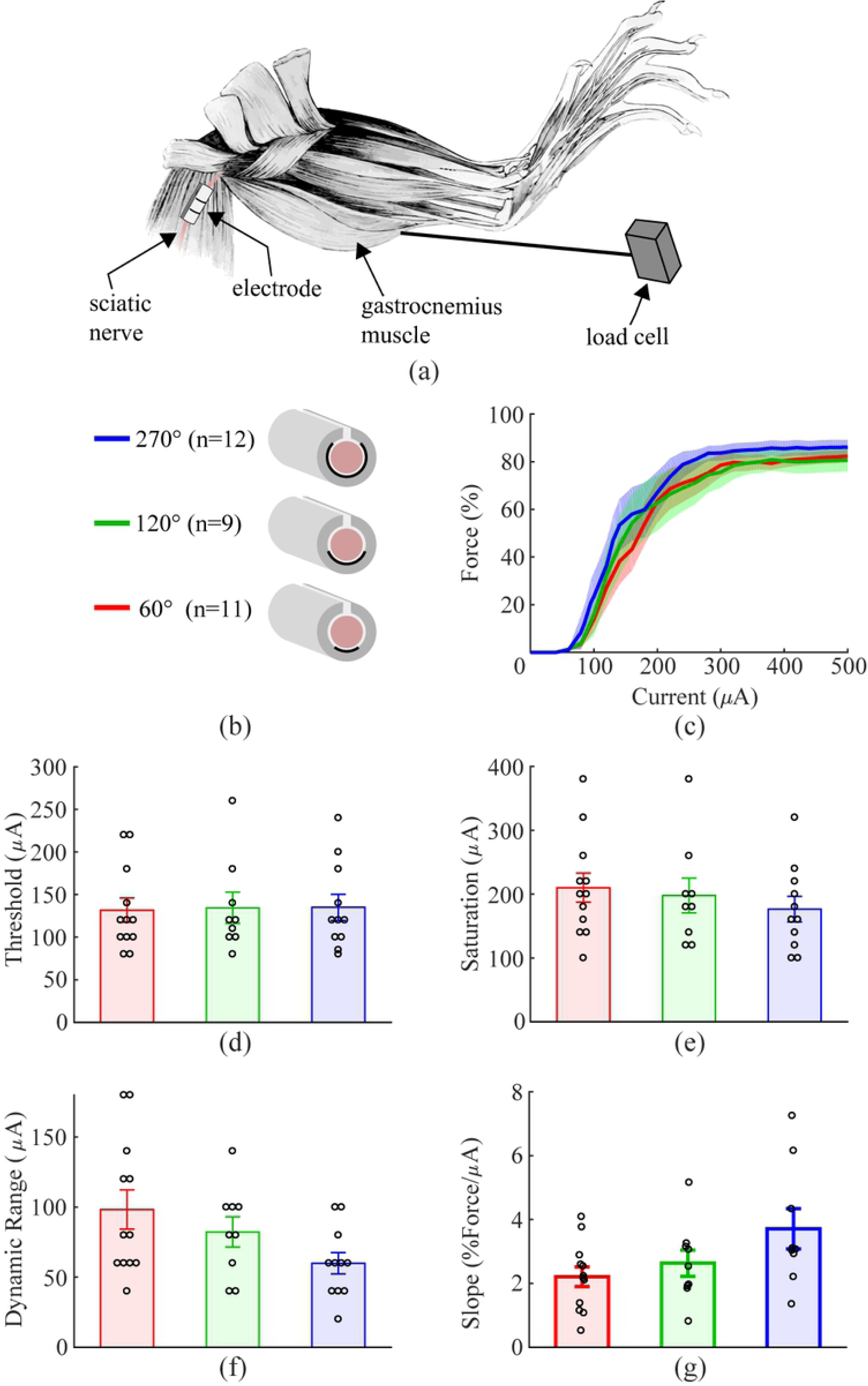
Approximately flat electrode does not reduce fiber recruitment in rat sciatic nerve. a) Schematic diagram of the experimental setup. b) Schematic diagram of the three cuff electrode designs tested on the rat sciatic nerve. c) Force generated as a function of stimulation intensity for each electrode design. All geometries result in similar recruitment. Shaded regions represent SEM. d-g) Thresholds, saturation currents, dynamic ranges, and slopes are similar for each electrode design. Data indicate mean ± SEM, and circles represent individual data.

### One-sided electrodes recruit more efficiently than circumferential electrodes in the rat vagus nerve

We next tested recruitment using the same 60° and 270° cuff electrodes on the rat vagus nerve, which has a smaller diameter and different fascicular organization.

#### Model

Modeling of the rat vagus showed an unexpected result: decreasing the angle of the electrodes improved recruitment by decreasing both the threshold and saturation current (Fig. 4a; 30°: *i*_5%_=101.6 µA, *i*_95%_=224.3 µA; 270°: *i*_5%_=262.0 µA, *i*_95%_=532.4 µA). To explain this result, three follow up tests were run. In the original model, the vagus nerve was positioned at the bottom of the cuff lumen very close to the contacts to match the experimental prep. The first two follow up tests changed the nerve’s position to be either in the middle of the cuff or on the opposite side from the contacts. When the nerve was in the middle of the cuff, changing the angle of coverage had no effect on recruitment (Fig. 4b; 30°: *i*_5%_=315.9 µA, *i*_95%_=679.4 µA; 270°: *i*_5%_=324.3 µA, *i*_95%_=674.0 µA). When the nerve was on the opposite side of the cuff, reducing angle of coverage increased both the threshold and saturation current (Fig. 4c; 30°: *i*_5%_=467.4 µA, *i*_95%_=1013.9 µA; 270°: *i*_5%_=357.9 µA, *i*_95%_=735.5 µA). The final follow up test varied the angle of coverage inside of a cuff which was properly sized for the rat vagus (0.44 mm inner diameter). In this case, varying the angle of coverage once again had minimal effect on recruitment (Fig. 4d; 30°: *i*_5%_=20.5 µA, *i*_95%_=42.0 µA; 270°: *i*_5%_=24.2 µA, *i*_95%_=47.2 µA). These data suggest that in a cuff that is significantly larger than the nerve, the vagus benefits from the increased current density near the contacts present with shorter angles of coverage without being affected by the decreased current density far from the contacts.

**Fig 4.**
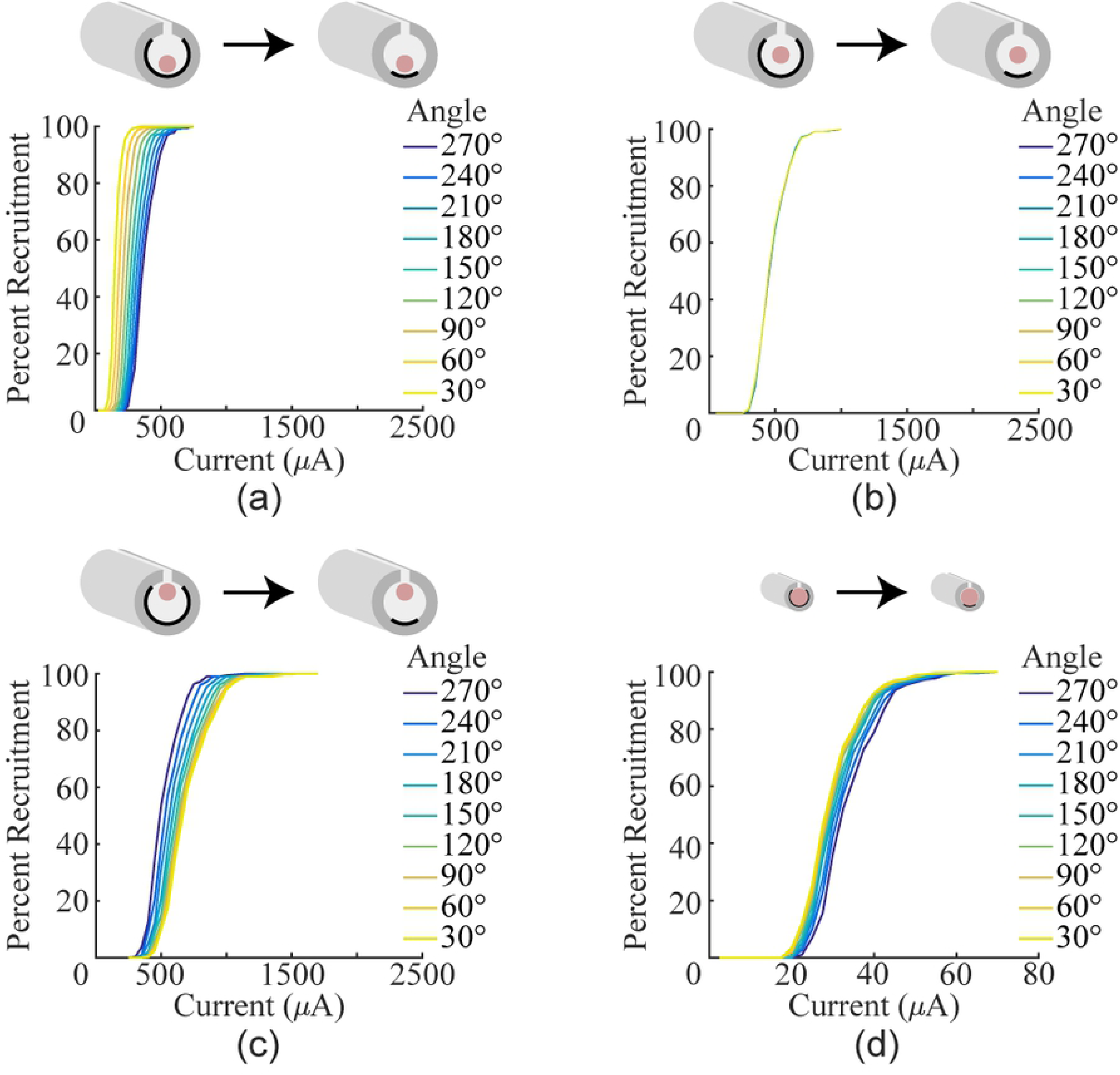
Reducing angle of coverage increases fiber recruitment in a model of the rat vagus nerve. a) Recruitment curves generated using a cuff with a 1 mm inner diameter, but with the nerve positioned at next to the contacts. Reducing the angle increases recruitment. b) Recruitment curves generated using a cuff with a 1 mm inner diameter, but with the nerve positioned in the middle of the cuff lumen. Reducing the angle has no effect. c) Recruitment curves generated using a cuff with a 1 mm inner diameter, but with the nerve on the opposite side of the cuff lumen from the contacts. Reducing the angle decreases recruitment. d) Recruitment curves generated by modeling cuff electrodes with various angles of completion around the rat vagus. Instead of a 1 mm inner diameter, the cuff diameter was set to 0.44 mm to keep the ratio of the cuff diameter to nerve the same as in the sciatic model. When the cuff is sized to fit the nerve, reducing the angle has little effect on fiber recruitment.

#### Empirical

We next sought to confirm these findings *in vivo*. To evaluate activation of vagus nerve fibers, we measured rapid stimulation-dependent reduction in oxygen saturation, a well-described biomarker of vagus nerve stimulation ascribed to activation of the Hering-Breuer reflex (32). Stimulation of vagal A-fibers, including the pulmonary stretch receptors, temporarily prevents inhalation and causes blood oxygen saturation to fall (Fig. 5a) (31). As a result, measurement of oxygen saturation provides a simple means to assess vagal A-fiber recruitment.

**Fig 5.**
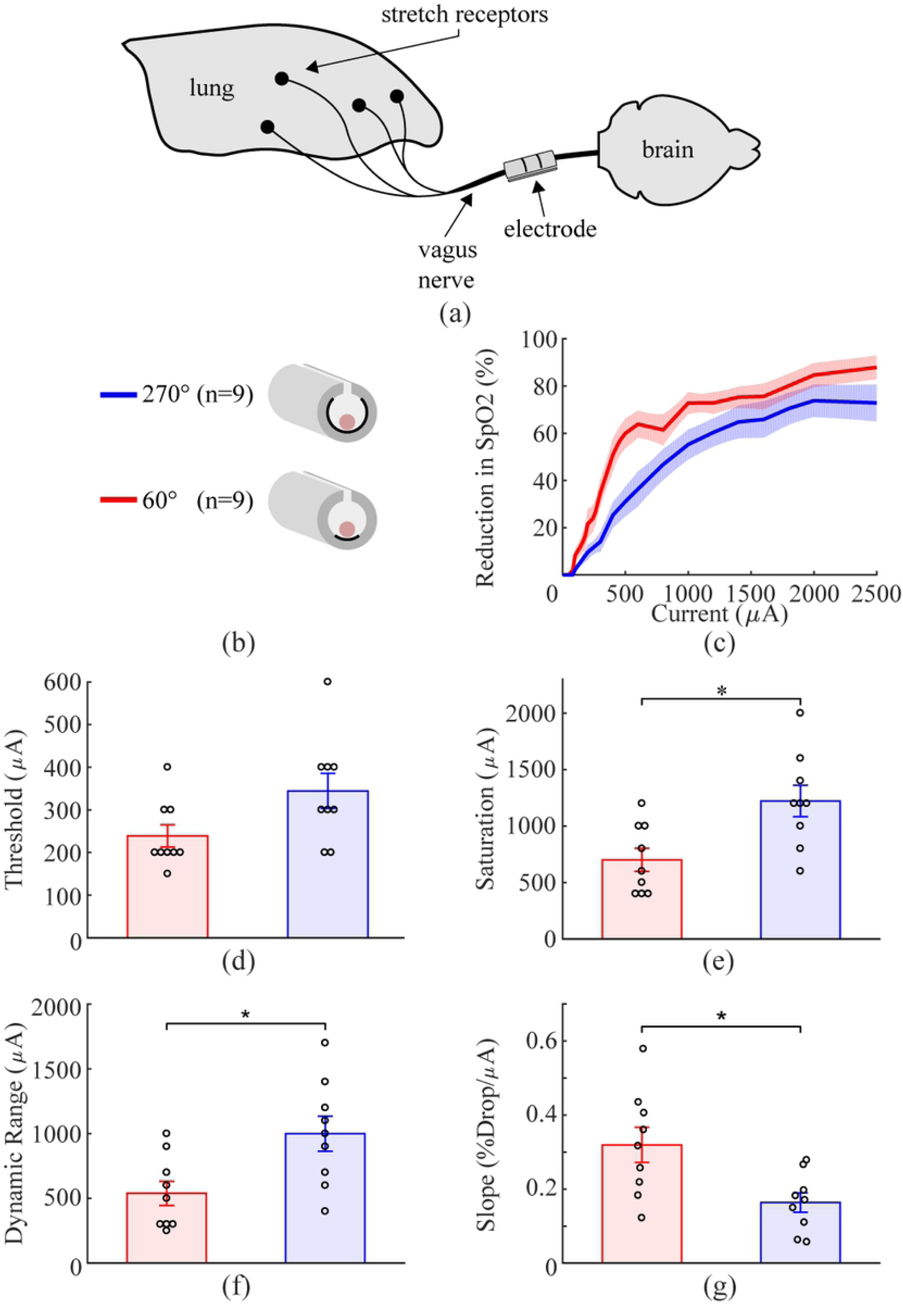
Reducing angle of coverage to approximate a flat electrode increases fiber recruitment in rat vagus nerve. a) Schematic diagram of the experimental setup. b) Schematic diagram of the two cuff electrode designs tested on the rat vagus nerve. c) Decreases in SpO2, a biomarker of vagal activation, as a function of stimulation intensity for each electrode design (y-axis is percent of maximum reduction). Similar to modeling results, the decreased angle of coverage generates more efficient nerve recruitment. d-g) Thresholds are similar for each design. The 60° electrodes displayed reduced saturation current, dynamic range, and increased slope compared to the 270° electrodes. Data indicate mean ± SEM, and circles represent individual data.

The 60° electrodes recruited fibers more effectively than the 270° electrodes, corroborating findings from the model. A trend toward reduced threshold was observed with the 60° electrode, although this failed to achieve statistical significance (Fig. 5d; 60°: 238.9 ± 26.1 µA, 270°: 344.4 ± 41.2 µA; two tailed paired t-test, p=0.0508). The 60° electrode displayed a significantly reduced saturation current, dynamic range, and increased slope compared to the 270° electrode (Saturation: Fig. 5e, 60°: 700 ± 102.7 µA, 270°: 1222 ± 139.2 µA, two tailed paired t-test, p=8.5×10^-3^; Dynamic Range: Fig. 5f, 60°: 538.9 ± 93.5 µA, 270°: 1000 ± 135.4 µA, two tailed paired t-test, p=0.014; Slope: Fig. 5g, 60°: 0.320 ± 0.047, 270°: 0.164 ± 0.026, two tailed paired t-test, p=0.0165). Both the model and empirical data demonstrate that the one-sided electrodes have a steeper recruitment curve and lower saturation current than circumferential electrodes.

### Flat and circumferential electrodes provide equivalent recruitment in rabbit sciatic nerve

The results presented above support the notion that flat electrodes provide at least as effective fiber recruitment as circumferential electrodes. However, whereas the 60° electrodes used in the above experiments contact only a single side of the nerve similar to a flat electrode, they are not truly flat and thus do not capture all the features of the geometry that may influence fiber recruitment. Therefore, we sought to confirm these results using a true flat electrode. The electrode was manufactured on a printed circuit board (PCB), encapsulated in glass, and inserted into a silicone sleeve that acted as an insulating cuff (Fig. 8c). These electrodes were tested on the rabbit sciatic nerve, which is an order of magnitude larger than the rat sciatic nerve (16,18).

#### Model

We performed modeling to evaluate recruitment using flat contacts and circumferential contacts. The cross-sectional area of the nerve and of the cuff lumen was matched between the circumferential and flat electrode models by increasing the inner diameter of the flat cuff by 8.67% and modifying the nerve shape to fit (Fig. 6) (23). Flat and circumferential designs had similar thresholds and saturation currents (Fig. 6; Flat: *i*_5%_=76.1 µA, *i*_95%_=278.6 µA; Circumferential: *i*_5%_=81.4 µA, *i*_95%_=253.1 µA). Next, flat and circumferential electrodes were compared in different ambient mediums with conductivities ranging from fat to saline. Decreasing the conductivity of the ambient medium increased recruitment, but the comparable recruitment observed with flat and circumferential contacts was consistent in all cases (Fig. 7a). Flat and circumferential electrodes were compared on nerves of varying size by increasing or decreasing the spatial scale of the original rabbit sciatic model. Larger nerves required more current to achieve similar levels of fiber recruitment, but once again, flat and circumferential electrodes were similar in all cases (Fig. 7b). A helical cuff electrode, like those used in clinical VNS applications, provides similar recruitment to the flat electrode design (Fig. S2). On a multi-fascicle nerve, flat and circumferential electrodes provided similar recruitment of the whole nerve despite each fascicle being recruited differently. These data suggest that flat electrodes recruit fibers similarly to currently used electrode designs.

**Fig 6.**
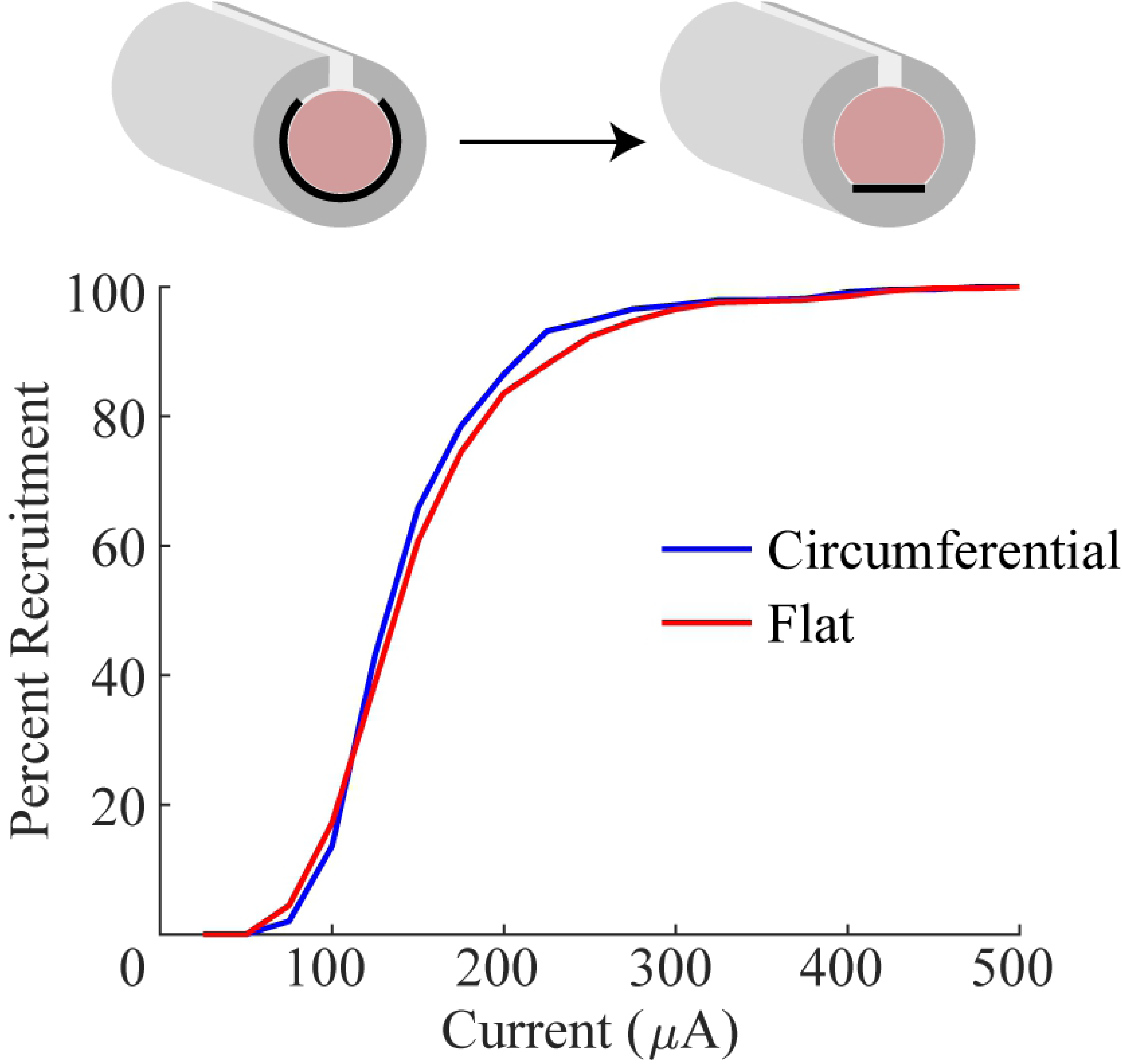
Flat and circumferential electrodes provide similar recruitment in a model of the rabbit sciatic nerve. Recruitment curves generated by modeling cuff electrodes around the rabbit sciatic nerve with either flat or circumferential contacts. Note the similarity in fiber recruitment.

**Fig 7.**
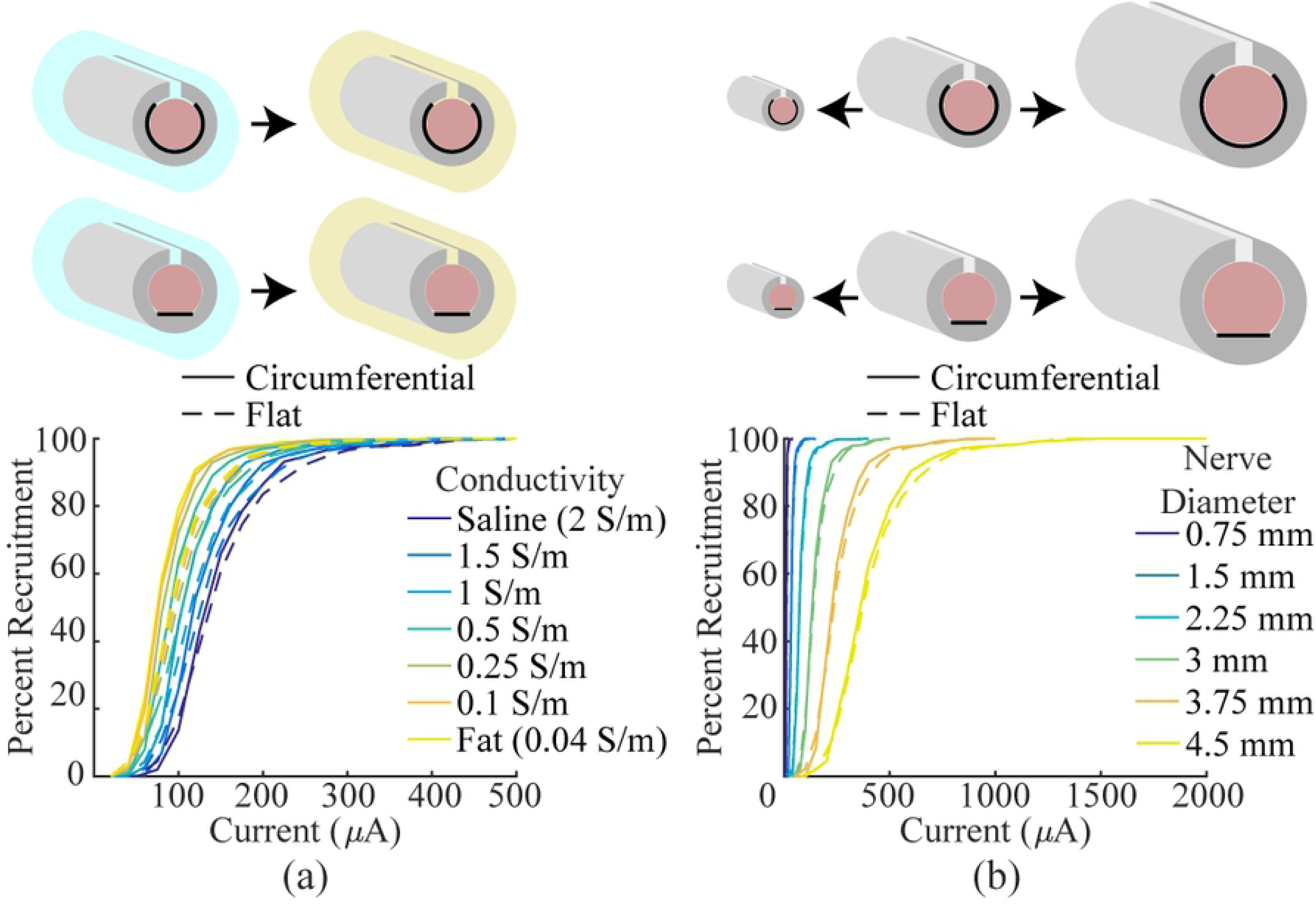
Models of flat and circumferential electrodes in various extracellular media and on various nerve sizes. a) Recruitment curves generated by modeling flat and circumferential electrodes in various ambient mediums. The conductivity of the ambient medium was varied from saline to fat. As expected, the extracellular medium influences recruitment efficiency, but recruitment is similar between the two electrode designs in all cases. b) Recruitment curves generated by modeling flat and circumferential electrodes on various diameter nerves. All features of the cuff electrode were kept proportional and scaled to match the nerve. In all cases, recruitment is similar between the two designs.

#### Empirical

To confirm modeling predictions, we evaluated nerve recruitment in the rabbit sciatic nerve by measuring the force of muscle contraction. No difference was found between the thresholds, saturation currents, or dynamic ranges (Threshold: Fig. 8d, Circumferential: 390.0 ± 14.8 µA, Flat: 351 ± 41.5 µA, two-sample F-test, F(5, 4)=9.397, p=0.0497, two tailed two-sample t-test with unequal variance: p=0.4167; Saturation: Fig. 8e, Circumferential: 514.0 ± 10.8 µA, Flat: 430.0 ± 46.0 µA, two-sample F-test, F(5, 4)=21.931, p=0.011, two tailed two-sample t-test with unequal variance: p=0.1301; Dynamic Range: Fig. 8f, Circumferential: 148.0 ± 21.5 µA, Flat: 111.7 ± 18.3 µA, two-sample F-test, F(5, 4)=0.8693, p=0.860, two tailed two-sample t-test with equal variance: p=0.228). These results confirm that flat electrodes recruit fibers equivalently to circumferential electrodes.

**Fig 8.**
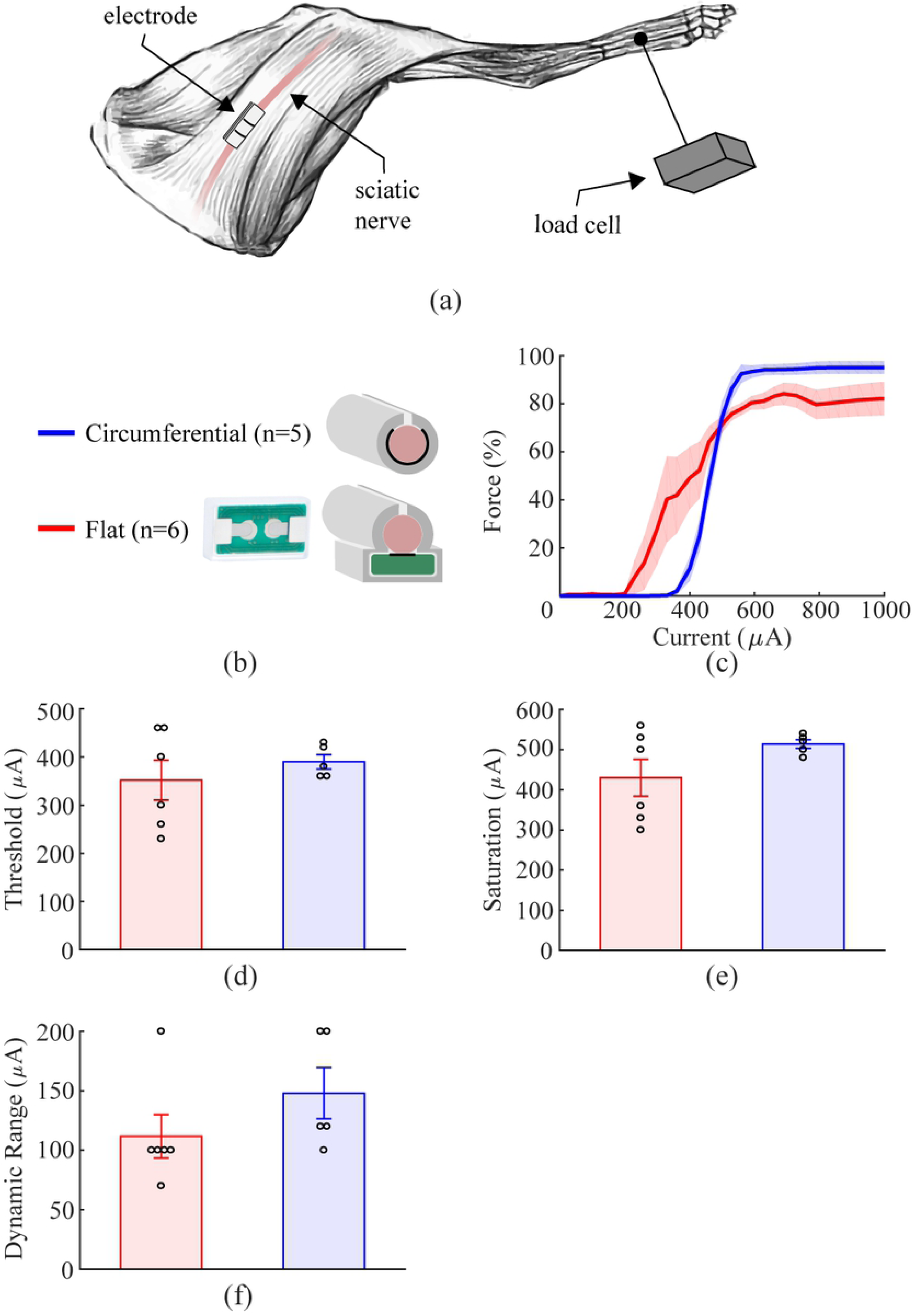
Flat and circumferential electrodes provide similar recruitment of rabbit sciatic nerve. a) Schematic diagram of the experimental setup. b) Schematic diagram of the two cuff electrode designs tested on the rabbit sciatic nerve. c) Force generated as a function of stimulation intensity for flat and circumferential electrodes. Both designs achieve efficient recruitment of the sciatic nerve, consistent with modeling predictions. d-f) Thresholds, saturation current, and dynamic range are similar for each electrode design. Data indicate mean ± SEM, and circles represent individual data.

## Discussion

In this study, we examined the viability of flat electrodes for nerve stimulation. Circumferential electrode contacts were compared to flat electrode contacts on multiple nerves and in multiple species. We find that in all cases tested, recruitment is either equivalent or the flat contacts have a steeper recruitment function and lower saturation current.

Flat electrodes that only contact a single side of the nerve could provide multiple advantages over the currently used circumferential designs which wrap around the majority of the circumference of the nerve. However, flat electrodes may require more current to achieve the same amount of fiber recruitment. On the rat sciatic nerve, both modeling and empirical data demonstrate that the reduced angle of coverage, approximating a flat electrode, provides comparable fiber recruitment to circumferential contacts across a wide range of current intensities. These results suggest that flat electrodes will recruit comparably to circumferential electrodes.

It is plausible that nerve diameter and fascicular organization could differentially affect recruitment with various electrode designs. Unexpectedly, the one-sided electrode contacts provided more efficient recruitment of the rat vagus nerve fibers than the circumferential contacts. There was once again no difference in the thresholds, but the 60° contacts had a steeper recruitment curve and lower saturation current, indicating more efficacious fiber recruitment. The modeling data confirmed these results, which can be ascribed to the relatively small size of the vagus nerve compared to the inner diameter of the insulating cuff. The diameter of the vagus nerve is around 0.4 mm, less than one half that of the sciatic, and thus the nerve occupies a substantially smaller cross-sectional area inside the cuff. Cuffs were always placed such that the vagus was resting at the bottom of the cuff and in the middle of the contacts. Additionally, injection current density was higher with the 60° electrodes given their reduced surface area compared to the 270° electrodes (41). Due to the small size of the nerve relative to the cuff, its position, and the increased current density near the contacts present with the 60° design, the current density within the nerve was higher with the smaller contact angle. Model results suggest that this is only true when the nerve is at the bottom of the cuff and the cuff is significantly larger than the nerve. If the nerve was moved to the opposite side of the cuff, far away from the contacts, the opposite relationship was seen (Fig. 4c), and if the cuff was sized appropriately for the vagus, the 60° contacts did not appear significantly different from the 270° contacts (Fig. 4d). Regardless, the modeling and empirical data support the notion that flat electrodes provide at least equivalent fiber recruitment. Whereas the 60° electrode contacts used in the rat experiments contact only a single side of the nerve similar to flat contacts, it is still possible that a true flat electrode would yield significantly less effective fiber recruitment. Thus, we tested nerve activation in the rabbit sciatic nerve using a true flat electrode manufactured on a PCB and compared recruitment to a standard circumferential electrode. Similar to the rat experiments, modeling and empirical testing revealed no substantial difference in fiber recruitment between the flat and circumferential electrode contacts. These results provide further evidence in an independent replicate that flat contacts stimulate as effectively as circumferential contacts. Furthermore, the devices used in these experiments were simple PCBs, which illustrates the convenience of using flat contacts.

For applications in which recruitment of all areas within the nerve is desired, flat electrodes appear to be a suitable alternative to the standard circumferential and helical electrodes. However, it is still unclear how each electrode design recruits individual fascicles within the nerve. Depending on the orientation of the nerve relative to the electrodes, the threshold for activation of any given fascicle will change (42). If there is any difference between electrode types, it is likely small as there were no significant differences in the empirical data, and both the vagus and the sciatic nerves have several fascicles, but this could explain the high variance in thresholds seen when testing flat electrodes on the rabbit sciatic nerve. The threshold for stimulation should be low if the target fascicle is near the electrode, but high if the fascicle is far from the electrode. A modified rabbit sciatic nerve with five fascicles was modeled with both flat and circumferential electrodes to investigate how each design is affected by the orientation of the nerve inside the cuff. The steepness of the recruitment curve for single fascicles was similar in all cases, but the variance in thresholds was greater with the flat electrodes (Fig. 9), which provides an explanation for the variance in thresholds present empirically. Fascicles on the opposite side of the nerve from the flat electrodes will have a higher threshold of activation, but the threshold is still comparable to circumferential electrodes, which suggests that clinical efficacy will not be reduced for applications in which whole nerve recruitment is desired.

**Fig 9.**
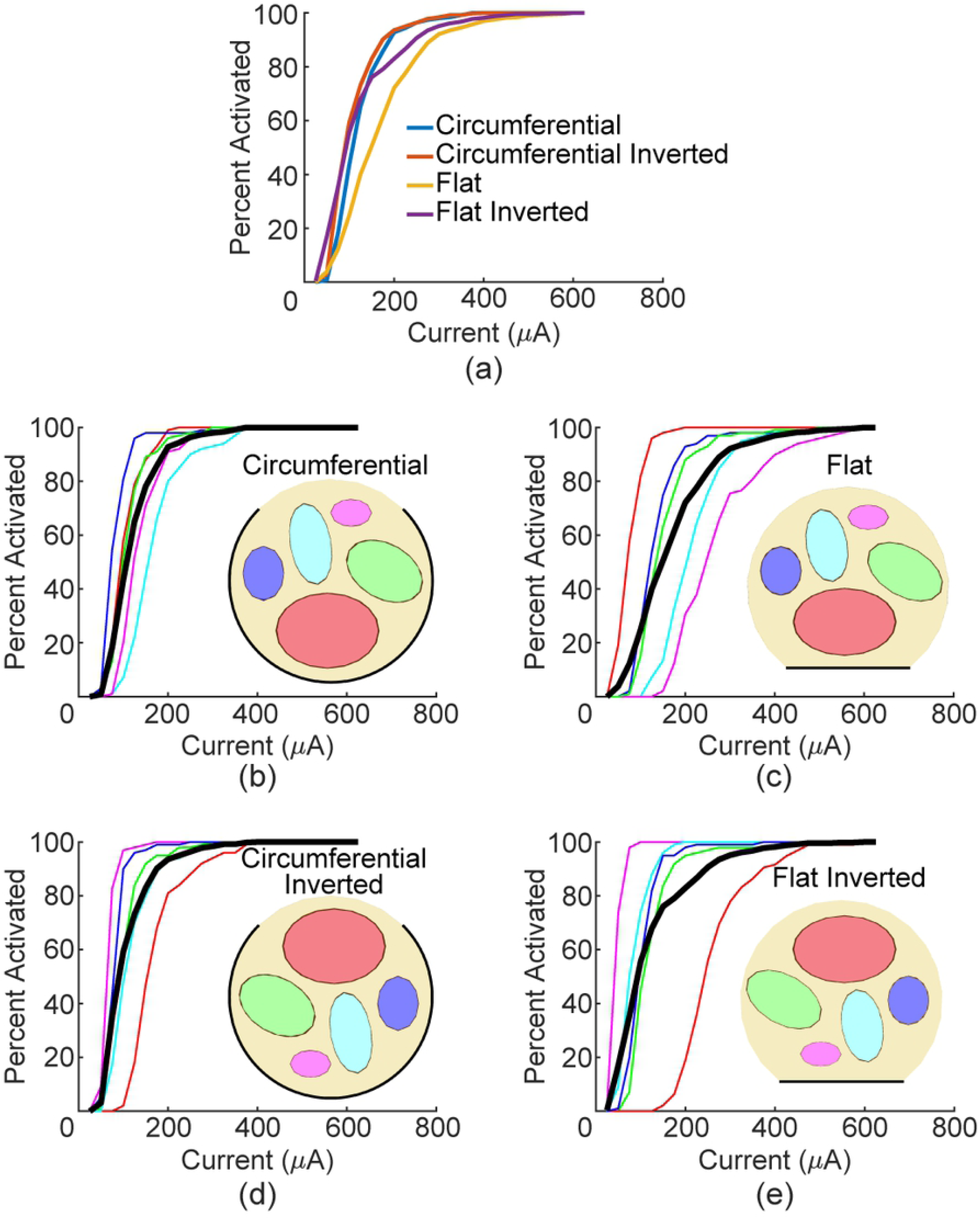
Flat electrodes result in greater threshold variability for individual fascicles, but similar recruitment of the whole nerve. a) Whole nerve recruitment curves for the four combinations modeled. b-e) Recruitment curves for each fascicle (line color corresponds to fascicle of same color) and whole nerve recruitment (thick black line).

All modeling and *in vivo* experiments in this study measured A-fiber recruitment, but many applications of nerve stimulation rely on B- and C-fibers as well (43). It is not guaranteed that flat and circumferential electrodes will recruit these other fiber types equivalently, but models of smaller diameter fibers suggest that the increase in fiber recruitment threshold would scale proportionally between the two electrode designs. Thus, while more current is required to activate smaller diameter fibers, we predict that the increase in current is likely to be similar comparing flat and circumferential designs. Further work validating this finding *in vivo* is warranted to determine if flat electrodes are viable for broader applications of nerve stimulation.

A major limitation of this study is the absence of empirical testing with chronically implanted electrodes. Many changes occur chronically that could result in reduced efficacy of flat electrodes such as glial scar formation, inflammation, and nerve damage (44). It is possible that some of these phenomena will affect flat electrodes differently than circumferential electrodes leading to electrode failure. Although chronic implants were not experimentally tested in this study, our modeling studies suggest that flat and circumferential electrodes provide comparable recruitment in a range of physiologically plausible extracellular media, which suggests that scar formation will not affect flat electrodes to a greater extent than it does circumferential electrodes. Future studies are needed to provide a direct empirical evaluation of the chronic efficacy of flat electrodes.

Additionally, there is a lack of data on larger diameter nerves (>3 mm) that would be found in humans. Our modeling studies suggest that flat and circumferential electrodes are equivalent on a range of clinically applicable nerve sizes. Moreover, comparison of recruitment in the rat sciatic and rabbit sciatic suggests that larger nerves require more current to achieve the same level of activation, but in both cases recruitment is comparable between flat and circumferential electrodes.

Our finding that larger nerves require more current to achieve similar levels of activation is intuitive. However, there is a strong body of literature showing that equivalent stimulation parameters can successfully activate rat and human nerves. This data is particularly compelling for the vagus nerve where both rats and humans exhibit enhanced memory as an inverted-U function of current intensity with the same peak (45–48). Our modeling efforts suggest a simple explanation for this surprising finding, which appears to lie in the use of tight-fitting stimulating electrodes for human studies and poorly fitting, oversized cuff electrodes for rat studies. When we modeled these configurations, we confirmed that identical VNS parameters can equivalently activate nerves of very different diameters under these conditions (Fig. 10). This is a novel result that could substantially impact both preclinical and clinical stimulation parameters. Follow up studies comparing small diameter nerves in animals to large diameter nerves in humans should be done to confirm this finding.

**Fig 10:**
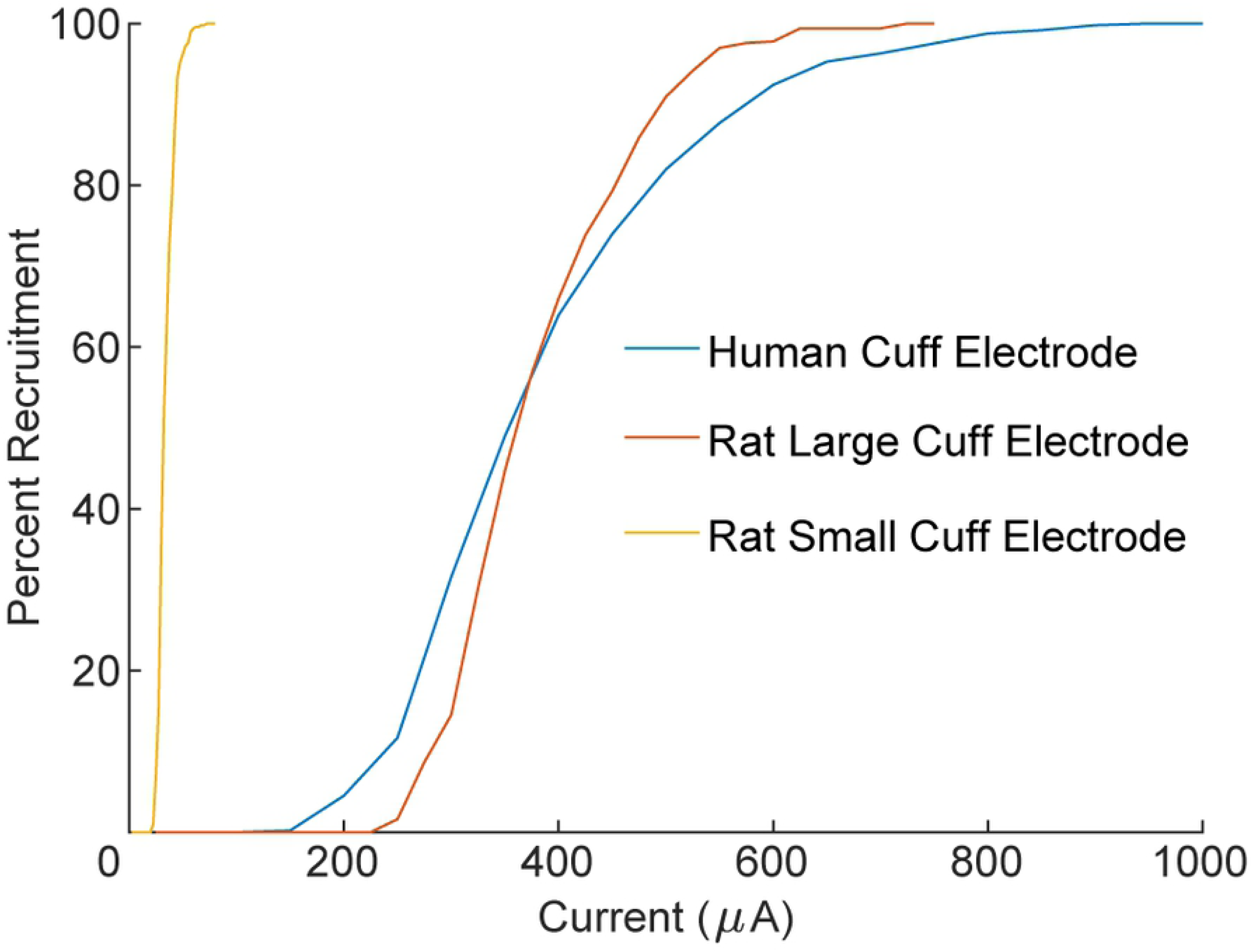
Recruitment of vagus nerve is similar in humans and rats due to cuff electrode design. Larger nerves require more current to recruit, but the therapeutic range of vagus nerve stimulation is similar in rats and humans (Fig. 7b). This phenomenon can be explained by the use of tight-fitting stimulating electrodes for human studies and poorly fitting, oversized cuff electrodes for rat studies. Cuff electrodes used in rats are significantly larger than the nerve which leads to inefficient recruitment and brings the two curves into alignment. If rat cuff electrodes were reduced in size, recruitment would be greatly increased. This is consistent with the importance of the ratio of cuff inner diameter to nerve diameter (Fig. 2a).

If a flat electrode design is used to stimulate the vagus nerve for epilepsy, it is not initially clear whether the stimulation parameters would be different from current clinical parameters, as a helical electrode design is used in clinical applications rather than a cuff electrode (49). We modeled the helical electrode design and compared vagus nerve recruitment to recruitment using a flat electrode design. The helical electrode and flat electrode demonstrated comparable recruitment. The narrow insulating structure used by the helical cuff allows some current to bypass the nerve, which increases the amount of stimulation needed compared to a complete cuff electrode (Fig. S2). The open architecture of the helical cuff is equivalent to having very little cuff overhang, which decreases recruitment compared to a full cuff (Fig. 2c).

These results provide a framework to guide the development of new electrode designs for nerve stimulation. The difference in fiber recruitment between flat and circumferential contacts is not likely to meaningfully influence the efficacy of nerve stimulation techniques, and flat contacts may provide advantages such as greater ease of implantation, substantially reduced cost of production, and greater resistance to mechanical failure. Future studies examining the effects of electrode size and geometry may provide further insights into design features to optimize recruitment for nerve stimulation therapies.

## Acknowledgements

We would like to thank Stuart Cogan, Joseph Pancrazio, Jonathan Riley, Nikki Simmons, and Brandon Tran for help with modeling of fibers in NEURON, equipment setup, physiological recordings, insightful discussions, figure creation, and manuscript preparation.

## References

1. Handforth A, DeGiorgio CM, Schachter SC, Uthman BM, Naritoku DK, Tecoma ES, et al. Vagus nerve stimulation therapy for partial-onset seizures. A randomized active-control trial. Neurology [Internet]. 1998;51(1):48–55. Available from: http://n.neurology.org/content/51/1/48

2. Kimberley TJ, Pierce D, Prudente CN, Francisco GE, Yozbatiran N, Smith P, et al. Vagus Nerve Stimulation Paired With Upper Limb Rehabilitation After Chronic Stroke. Stroke [Internet]. 2018;49:1–4. Available from: https://www.ahajournals.org/doi/abs/10.1161/STROKEAHA.118.022279

3. De Ridder D, Vanneste S, Engineer ND, Kilgard MP. Safety and efficacy of vagus nerve stimulation paired with tones for the treatment of tinnitus: A case series. Neuromodulation Technol Neural Interface. 2014;17(2):170–9.

4. Tassorelli C, Grazzi L, de Tommaso M, Pierangeli G, Martelletti P, Rainero I, et al. Noninvasive vagus nerve stimulation as acute therapy for migraine. Neurology [Internet]. 2018;91(4):364–73. Available from: http://www.neurology.org/lookup/doi/10.1212/WNL.0000000000005857

5. MacDiarmid SA, Peters KM, Shobeiri SA, Wooldridge LS, Rovner ES, Leong FC, et al. Long-Term Durability of Percutaneous Tibial Nerve Stimulation for the Treatment of Overactive Bladder. J Urol [Internet]. 2010;183(1):234–40. Available from: http://dx.doi.org/10.1016/j.juro.2009.08.160

6. Kamm MA, Dudding TC, Melenhorst J, Jarrett M, Wang Z, Buntzen S, et al. Sacral nerve stimulation for intractable constipation. Gut. 2010;59(3):333–40.

7. Saper JR, Dodick DW, Silberstein SD, McCarville S, Sun M, Goadsby PJ. Occipital nerve stimulation for the treatment of intractable chronic migraine headache: ONSTIM feasibility study. Cephalalgia. 2011;31(3):271–85.

8. Eastwood PR, Barnes M, Walsh JH, Maddison KJ, Hee G, Schwartz AR, et al. Treating obstructive sleep apnea with hypoglossal nerve stimulation. Sleep [Internet]. 2011;34(11):1479– 86. Available from: http://www.ncbi.nlm.nih.gov/pubmed/22043118

9. Birmingham K, Gradinaru V, Anikeeva P, Grill WM, Pikov V, Mclaughlin B, et al. Bioelectronic medicines: a research roadmap. 2014 [cited 2018 Jul 23]; Available from: http://dx.doi.org/10.1038/

10. Tan D, Schiefer M, Keith MW, Anderson R, Tyler DJ. Stability and selectivity of a chronic, multi-contact cuff electrode for sensory stimulation in a human amputee. Int IEEE/EMBS Conf Neural Eng NER. 2013;859–62.

11. Boretius T, Badia J, Pascual-Font A, Schuettler M, Navarro X, Yoshida K, et al. A transverse intrafascicular multichannel electrode (TIME) to interface with the peripheral nerve. Biosens Bioelectron [Internet]. 2010;26(1):62–9. Available from: http://dx.doi.org/10.1016/j.bios.2010.05.010

12. Badia J, Boretius T, Andreu D, Azevedo-Coste C, Stieglitz T, Navarro X. Comparative analysis of transverse intrafascicular multichannel, longitudinal intrafascicular and multipolar cuff electrodes for the selective stimulation of nerve fascicles. J Neural Eng. 2011;8(3).

13. Kahlow H, Olivecrona M. Complications of vagal nerve stimulation for drug-resistant epilepsy: A single center longitudinal study of 143 patients. Seizure [Internet]. 2013;22(10):827–33. Available from: http://dx.doi.org/10.1016/j.seizure.2013.06.011

14. Mourdoukoutas AP, Truong DQ, Adair DK, Simon BJ, Bikson M. High-Resolution Multi-Scale Computational Model for Non-Invasive Cervical Vagus Nerve Stimulation. Neuromodulation Technol Neural Interface [Internet]. 2017;2017. Available from: http://doi.wiley.com/10.1111/ner.12706

15. Helmers SL, Begnaud J, Cowley A, Corwin HM, Edwards JC, Holder DL, et al. Application of a computational model of vagus nerve stimulation. Acta Neurol Scand. 2012;126(5):336–43.

16. Shen J, Wang H-Q, Zhou C-P, Liang B-L. MAGNETIC RESONANCE MICRONEUROGRAPHY OF RABBIT SCIATIC NERVE ON A 1.5-T CLINICAL MR SYSTEM CORRELATED WITH GROSS ANATOMY. Microsurgery. 2010;28(1):32–6.

17. Woodbury JW, Woodbury DM. Vagal Stimulation Reduces the Severity of Maximal Electroshock Seizures in Intact Rats: Use of a Cuff Electrode for Stimulating and Recording. Pacing Clin Electrophysiol. 1991;14(1):94–107.

18. Varejão ASP, Cabrita AM, Meek MF, Bulas-Cruz J, Melo-Pinto P, Raimondo S, et al. Functional and Morphological Assessment of a Standardized Rat Sciatic Nerve Crush Injury with a Non-Serrated Clamp. J Neurotrauma [Internet]. 2004;21(11):1652–70. Available from: http://www.liebertonline.com/doi/abs/10.1089/neu.2004.21.1652

19. Grinberg Y, Schiefer MA, Tyler DJ, Gustafson KJ. Fascicular perineurium thickness, size, and position affect model predictions of neural excitation. IEEE Trans Neural Syst Rehabil Eng. 2008;16(6):572–81.

20. Yoo PB, Lubock NB, Hincapie JG, Ruble SB, Hamann JJ, Grill WM. High-resolution measurement of electrically-evoked vagus nerve activity in the anesthetized dog. J Neural Eng. 2013;10(2).

21. Somann JP, Albors GO, Neihouser K V., Lu KH, Liu Z, Ward MP, et al. Chronic cuffing of cervical vagus nerve inhibits efferent fiber integrity in rat model. J Neural Eng. 2018;15(3).

22. Islam MS, Oliveira MC, Wang Y, Henry FP, Randolph MA, Park BH, et al. Extracting structural features of rat sciatic nerve using polarization-sensitive spectral domain optical coherence tomography. J Biomed Opt [Internet]. 2012;17(5):056012. Available from: http://biomedicaloptics.spiedigitallibrary.org/article.aspx?doi=10.1117/1.JBO.17.5.056012

23. Tyler DJ, Durand DM. Functionally selective peripheral nerve stimulation with a flat interface nerve electrode. IEEE Trans Neural Syst Rehabil Eng. 2002;10(4):294–303.

24. Veltink PH, Van Veen BK, Struijk JJ, Holsheimer J, Boom HBK. A Modeling Study of Nerve Fascicle Stimulation. IEEE Trans Biomed Eng. 1989;36(7):683–92.

25. Goodall E V., Kosterman LM, Holsheimer J, Struijk JJ. Modeling Study of Activation and Propagation Delays During Stimulation of Peripheral Nerve Fibers with a Tripolar Cuff Electrode. IEEE Trans Rehabil Eng. 1995;3(3):272–82.

26. Frieswijk TA, Smit JPA, Rutten WLC, Boom HBK. Force-current relationships in intraneural stimulation: Role of extraneural medium and motor fibre clustering. Med Biol Eng Comput. 1998;36(4):422–9.

27. Arle JE, Carlson KW, Mei L. Investigation of mechanisms of vagus nerve stimulation for seizure using finite element modeling. Epilepsy Res [Internet]. 2016;126:109–18. Available from: http://dx.doi.org/10.1016/j.eplepsyres.2016.07.009

28. McIntyre CC, Richardson AG, Grill WM. Modeling the Excitability of Mammalian Nerve Fibers: Influence of Afterpotentials on the Recovery Cycle. J Neurophysiol [Internet]. 2002;87(2):995– 1006. Available from: http://jn.physiology.org/lookup/doi/10.1152/jn.00353.2001

29. Ikeda M, Oka Y. The relationship between nerve conduction velocity and fiber morphology during peripheral nerve regeneration. Brain Behav. 2012;2(4):382–90.

30. Germana G, Muglia U, Santoro M, Abbate F, Laura R, Gugliotta MA, et al. Morphometric analysis of sciatic nerve and its main branches in the rabbit. Biol Struct Morphog [Internet]. 1992;4(1):11–5. Available from: http://www.ncbi.nlm.nih.gov/entrez/query.fcgi?cmd=Retrieve&db=PubMed&dopt=Citation&list_uids=1420593

31. Chang R, Strochlic D, Williams E, Umans B, Liberles S. Vagal Sensory Neuron Subtypes that Differentially Control Breathing. Cell. 2015;161(3):622–33.

32. McAllen RM, Shafton AD, Bratton BO, Trevaks D, Furness JB. Calibration of thresholds for functional engagement of vagal A, B and C fiber groups in vivo. Bioelectron Med [Internet]. 2018;1(1):21–7. Available from: http://www.futuremedicine.com/doi/10.2217/bem-2017-0001

33. Gasser HS, Grundfest H. AXON DIAMETERS IN RELATION TO THE SPIKE DIMENSIONS AND THE CONDUCTION VELOCITY IN MAMMALIAN A FIBERS. Am J Physiol. 1939;127(2):393–414.

34. Hursh JB. Conduction Velocity and Diameter of Nerve Fibers. Am J Physiol Content. 1939;127(1):131–9.

35. Rios MU, Bucksot JE, Rahebi KC, Engineer CT, Michael P. Protocol for Construction of Rat Nerve Stimulation Cuff Electrodes. Methods Protoc. 2019;2(19):1–27.

36. Sivaji V. Wireless Devices for Peripheral Nerve Stimulation and Recording. University of Texas at Dallas; 2018.

37. Branner A, Stein RB, Fernandez E, Aoyagi Y, Normann RA. Long-Term Stimulation and Recording with a Penetrating Microelectrode Array in Cat Sciatic Nerve. IEEE Trans Biomed Eng. 2004;51(1):146–57.

38. Borland MS, Vrana WA, Moreno NA, Fogarty EA, Buell EP, Sharma P, et al. Cortical Map Plasticity as a Function of Vagus Nerve Stimulation Intensity. Brain Stimul [Internet]. 2016;9(1):117–23. Available from: http://dx.doi.org/10.1016/j.brs.2015.08.018

39. Ganzer PD, Darrow MJ, Meyers EC, Solorzano BR, Ruiz AD, Robertson NM, et al. Closed-loop neuromodulation restores network connectivity and motor control after spinal cord injury. Elife [Internet]. 2018;7:1–19. Available from: https://elifesciences.org/articles/32058

40. Loerwald KW, Borland MS, Rennaker RL, Hays SA, Kilgard MP. The interaction of pulse width and current intensity on the extent of cortical plasticity evoked by vagus nerve stimulation. Brain Stimul [Internet]. 2017;11(2):271–7. Available from: https://doi.org/10.1016/j.brs.2017.11.007

41. Cogan SF. Neural Stimulation and Recording Electrodes. Annu Rev Biomed Eng [Internet]. 2008 [cited 2018 Jul 30];10:275–309. Available from: www.annualreviews.org

42. Grill WM, Mortimer JT. Quantification of recruitment properties of multiple contact cuff electrodes. IEEE Trans Rehabil Eng. 1996;4(2):49–62.

43. Qing KY, Wasilczuk KM, Ward MP, Phillips EH, Vlachos PP, Goergen CJ, et al. B fibers are the best predictors of cardiac activity during Vagus nerve stimulation Qing, vagal B fiber activation and cardiac effects. 2018;1–11.

44. Cheung KC. Implantable microscale neural interfaces. 2007;(January):923–38.

45. Clark KB, Krahl SE, Smith DC, Jensen RA. Post-training unilateral vagal stimulation enhances retention performance in the rat. Vol. 63, Neurobiology of Learning and Memory. 1995. p. 213–6.

46. Clark KB, Smith DC, Hassert DL, Browning RA, Naritoku DK, Jensen RA. Posttraining Electrical Stimulation of Vagal Afferents with Concomitant Vagal Efferent Inactivation Enhances Memory Storage Processes in the Rat [Internet]. 1998 [cited 2018 Aug 1]. Available from: https://ac.els-cdn.com/S1074742798938631/1-s2.0-S1074742798938631-main.pdf?_tid=4779abf5-6708-44af-8a1f-2270db3602dc&acdnat=1533159129_91ad974012e12b5efe81cc4cc5798a6b

47. Clark KB, Naritoku DK, Smith DC, Browning RA, Jensen RA. Enhanced recognition memory following vagus nerve stimulation in human subjects. Nat Neurosci [Internet]. 1999 [cited 2018 Aug 1];2(1):94–8. Available from: http://neurosci.nature.com

48. Zuo Y, Smith DC, Jensen RA. Vagus nerve stimulation potentiates hippocampal LTP in freely-moving rats. Physiol Behav [Internet]. 2007 [cited 2018 Aug 15];90:583–9. Available from: https://ac.els-cdn.com/S0031938406004963/1-s2.0-S0031938406004963-main.pdf?_tid=9a134a03-79fd-4530-8d64-e33eecd8ddf7&acdnat=1534354220_560f2f94ec5032890fbe8a759e9cf349

49. Ben-Menachem E, Mañon-Espaillat R, Ristanovic R, Wilder BJ, Stefan H, Mirza W, et al. Vagus Nerve Stimulation for Treatment of Partial Seizures: 1. A Controlled Study of Effect on Seizures. Epilepsia. 1994;35(3):616–26.

